# Mating activates neuroendocrine pathways signaling hunger in *Drosophila* females

**DOI:** 10.1101/2022.10.19.512959

**Authors:** Meghan Laturney, Gabriella R. Sterne, Kristin Scott

## Abstract

Mated females reallocate resources to offspring production, causing changes in nutritional requirements and challenges to energy homeostasis. Although observed in most species, the neural and endocrine mechanisms that regulate the nutritional needs of mated females are not well understood. Here, we investigate the neural circuitry that regulates sugar appetite in mated *Drosophila melanogaster* females. During copulation, a male-derived sex peptide is transferred to females, silencing the mating status circuit to elicit many postmating behavioral changes^1-3^. We find that increased sucrose consumption is a postmated female behavior and show that it is mediated by the mating status circuit. We discovered that sexually dimorphic insulin receptor (Lgr3) neurons integrate mating status and nutritional state signals to adjust sucrose consumption. Lgr3+ cells receive inhibitory input from the mating status circuit via female specific pCd-2 neurons. In mated females, the inhibition of Lgr3 cells from pCd-2 is attenuated, transforming the mated signal into a long-term hunger signal that promotes sugar intake. Our results thus demonstrate that the mating circuit alters nutrient sensing centers in females to promote sugar consumption, providing a mechanism to increase intake in anticipation of the energetic costs associated with reproduction.

## Main

Female mating status impacts feeding decisions across species. In *Drosophila*, mated females increase total food intake^4^ and develop specific appetites for protein and salt^5-7^. Sugar, another macronutrient, is vitally important for flies and is their main energy source^8^. In mated females, increased activity^9^ results in elevated metabolic rates^10^ driving the need for additional calories. Moreover, females pack eggs with lipids^11^ synthesized from dietary carbohydrates^10^. Together, this predicts that mated females require significantly more sugar than virgins. However, as excessive sugar negatively impacts fly health^12^, sugar intake must be tightly regulated. Although the sugar to protein ratio influences reproductive output and longevity^8^, the absolute levels of sugar intake of mated females remain untested.

In *Drosophila*, mated related changes in females are orchestrated by sex peptide (SP), a 36-amino acid peptide that is produced in the male seminal fluid and transferred to females during copulation^1,2^. SP binds to its receptor (SPR)^13^ which is expressed in sensory neurons (SPSNs) in the uterus^14,15^. SPSNs convey mating status to sex peptide abdominal ganglia (SAG) neurons that ascend to the central brain^3^ and synapse onto pC1 neurons^16^. This female-specific SPSN-SAG-pC1 circuit is active in virgin females and silenced after mating^3,16^. Reception of SP and silencing of this circuit initiates the long-term postmating response in females, which includes changes in egg laying and sexual receptivity^3,16,17^. The second-order SAG neurons also drive changes in salt and protein appetites^7^, demonstrating that the mating status circuit alters food consumption in mated females. However, how the mating circuit influences nutritional state circuits or feeding circuits remains a central question. Here, we use automated behavioral assays for measuring consumption, powerful genetic tools, connectomics, and functional imaging approaches to examine neural mechanisms for appetite changes in mated females.

To investigate the impact of female mating status on sugar intake, we monitored individual feeding bouts over time using a high-throughput, automated feeding platform (FLIC^18^, Fig. 1a). We compared consumption of virgins and two types of mated females (1-hr or 72-hr postmated). We found that 72-hr postmated females consumed for a significantly longer duration on sucrose solution than virgins or 1-hr postmated females (Fig. 1b). To further evaluate the onset of increased sucrose consumption, we monitored consumption of virgin, 24-hr postmated females, and 72-hr psotmated females and established a postmated phenotype in both mated groups (Fig. 1c). These studies demonstrate that mated females consume more sugar than virgins and that changes in sucrose consumption manifest 6-24hr after copulation.

**Fig. 1.**
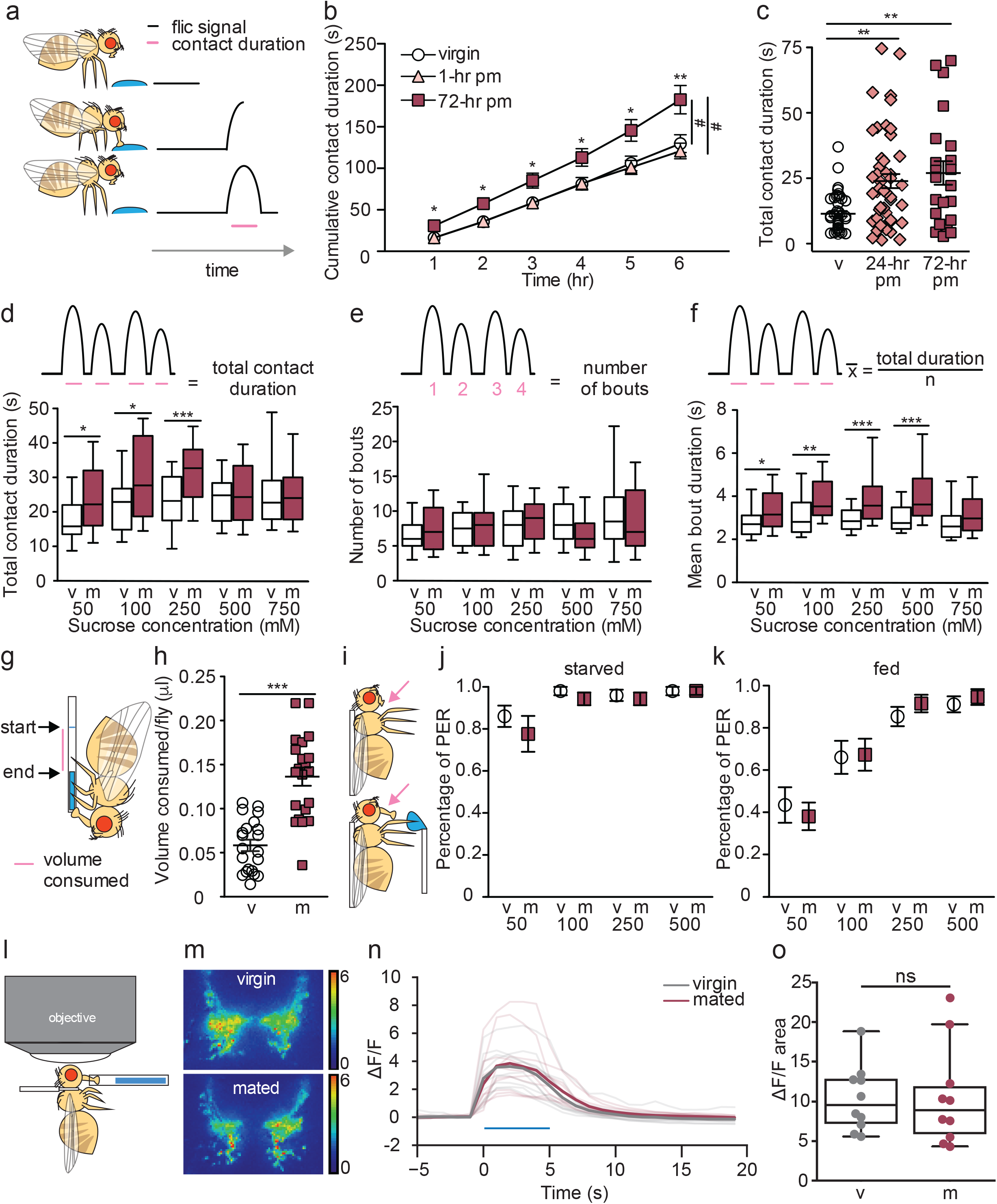
Mated females consume more sucrose than virgins by elongating feeding events. **a**, Schematic of fly feeding behavior (left) and the FLIC signal that monitors food contact over time (right). **b**, Cumulative drinking time in FLIC of virgin (white circles, n=34), 1-hr postmated (pm) (light pink triangles, n=30), and 72-hr pm (dark pink squares, n=28) females. Line graph shows mean and s.e.m. over time. Kruskal-Wallis was used to compare groups at hour intervals, Dunn’s posthoc. **c**, Total drinking time in FLIC in 20-min assay of virgin (v; n=47), 24-hr pm (n=47), and 72-hr pm (n=22) females, Kruskal-Wallis, Dunn’s posthoc. **d**-**f**, Schematic and plots of feeding behavior of virgin (v) and mated (m) females presented with sucrose of varying concentrations in FLIC 20-min assay (n=33-36) examining total drinking time (**d**), number of drinking bouts (**e**), and average bout duration (**f**). Box plot show first to third quartile, whiskers span 10-90 percentile, Mann Whitney. **g**, Schematic of female drinking from capillary containing sucrose solution in CAFE assay. **h**, Total volume consumed in CAFE assay per fly in 24 hours by virgin (v, n=21) and mated females (m, n=21), unpaired t-test. **i**, Schematic of proboscis extension response (PER) assay. **j**-**k**, Percentage of proboscis extensions observed per fly upon three presentations of sucrose of indicated concentration in virgin (v) and mated (m) females in starved (n=17-18, **j**) and fed state (n=16, **k**), Mann Whitney. **l**, Schematic of in vivo calcium imaging experiment monitoring taste-induced activity in gustatory axons. **m**-**o**, Changes in GCaMP6s signal in virgin (grey) and mated (dark pink) females, shown as representative fluorescent changes in single brains (**m**), changes over time (**n**), and maximum changes (**o**), n=10. Blue bar represents proboscis sucrose stimulation, Wilcoxen Rank test. *p<0.05, **p<0.01, ***p<0.001, ns = not significant. Scatter plot show mean +/-s.e.m. (**c**,**h**,**o**). Column graph show mean +/-s.e.m. (**j**,**k**). For full genotypes, see Supplementary Data Table 1.

To explore differences in feeding dynamics, virgin and mated females were allowed to feed on different sucrose concentrations and the number and length of feeding bouts were examined. Mated females consumed for longer durations than virgins for all sucrose concentrations below 500mM (Fig. 1d). Investigation into the microstructure of feeding revealed that the postmated increase in sucrose consumption is due to an elongation in feeding bout duration rather than an increased number of bouts (Fig. 1e,f). Thus, like hungry flies^19^, mated females consume more by engaging in longer feeding times rather than by initiating more feeding events.

We tested two predictions suggested by the mating-induced changes in sugar feeding dynamics. First, longer overall feeding durations suggest that a greater amount of sugar is consumed. We directly tested this using a capillary feeder (CAFE^20^, Fig. 1g) and found that mated females consume significantly more sucrose than virgins (Fig. 1h). Second, virgin and mated females execute a similar number of feeding bouts, suggesting that females do not differ in the probability of initiating a feeding event. To directly test this, we monitored proboscis extension upon sugar taste detection (Fig. 1i) and found that virgin and mated females have similar feeding initiation propensities (Fig. 1j,k). These results further corroborate that increased sugar consumption in mated females is due to longer meal duration rather than more frequent feeding bouts.

Hunger increases the sensitivity of gustatory neurons to sugar^21^. To test whether changes in sugar sensory sensitivity underlie increased sucrose consumption in mated females, we monitored taste-induced neural activity in sugar gustatory neurons in mated and virgin females with the GCaMP6s calcium indicator (Fig. 1l). Stimulation of the fly proboscis with sugar elicited similar responses in gustatory axons regardless of mating status (Fig. 1m-o), demonstrating that gustatory sensitivity is not altered by mating and not likely to contribute to changes in feeding.

As mating increases egg production, we reasoned that the increase in sucrose consumption could be driven by a need-dependent mechanism. In this scenario, mating induces egg production, depleting energy stores and driving a need for sucrose consumption to restore homeostasis. To test this, we quantified sucrose consumption of virgin and mated eggless females with both FLIC and CAFE (Fig. 2a and Extended Data Fig.1a, respectively). We found that despite a lack of egg production^7,22,23^, mated females consumed more sucrose than virgins, demonstrating that the change in postmated sucrose feeding is not driven by egg production itself. Instead, this result suggests that sucrose feeding changes are anticipatory in nature and a consequence of a need-independent mechanism.

**Fig. 2.**
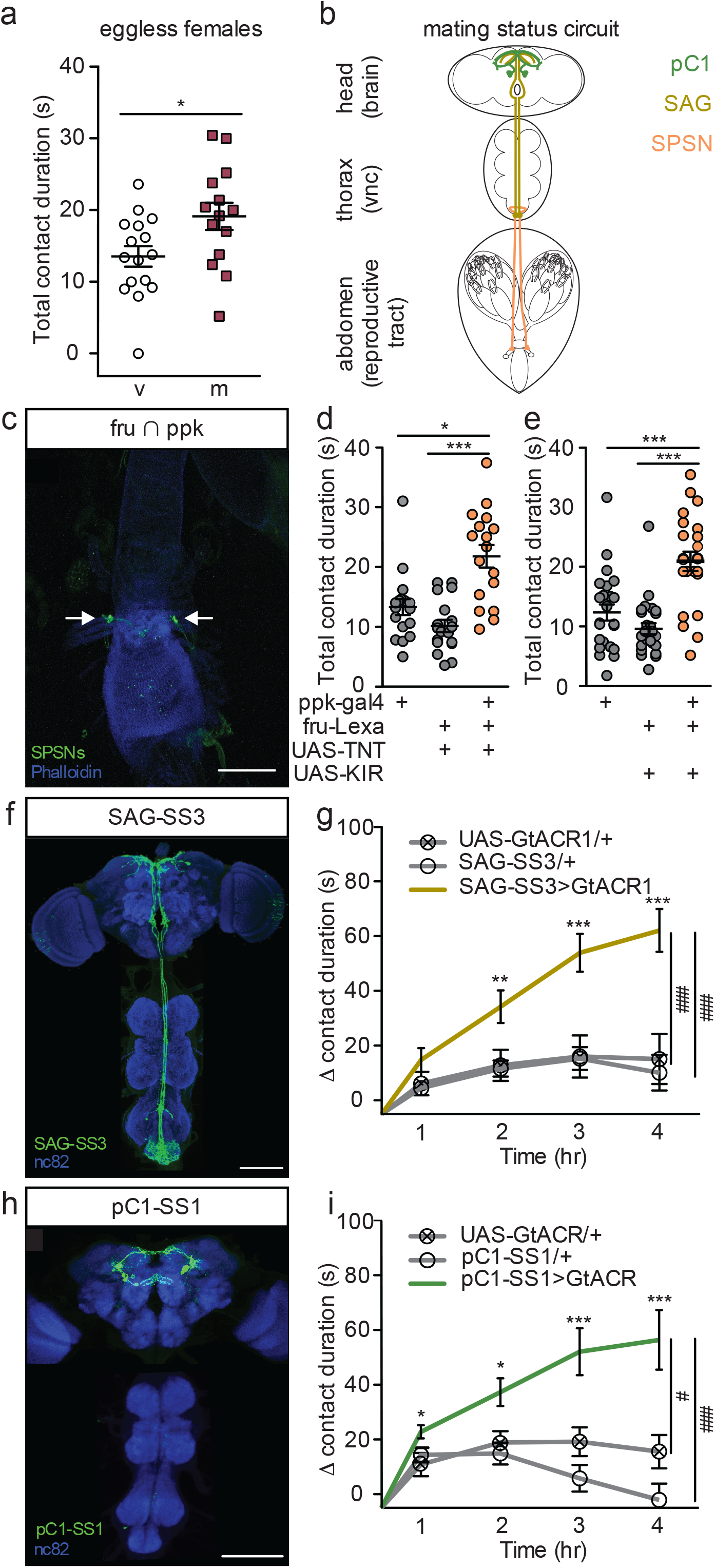
Postmated increases in sucrose consumption are induced by the canonical mating status circuit and are independent of egg-production. **a**, Total drinking time in FLIC in 20-min assay of eggless virgin (v, n=16) and mated (m, n=15) females, Mann Whit-ney. **b**, Schematic of female with the mating circuit, showing first order (SPSN, pink-orange), second order (SAG, dark yellow) and third order (pC1, green) neurons. **c**, Confocal image of uterus stained to reveal SPSNs (UAS-CD8:GFP, green, arrow-head indicating cell bodies) and muscle tissue (phalloidin, blue). **d**-**e**, Total drinking time in FLIC in 20-min assay of virgin females of indicated genotype (n=17-28), Kruskal-Wallis, Dunn’s posthoc. **f**,**h** Confocal images of brain (top) and ventral nerve cord (bottom) of SAG-SS3 (**f**) and pC1-SS1 (**h**) females, stained to reveal neural projection pattern (UAS-mVenus, green) and all synapses (nc82, blue). **g**-**i**, Drinking time of females exposed to light normalized to dark controls of indicated genotype in FLIC over 4-hrs (each data point is the drinking time of an individual in the light condition minus the daily average drinking time of all females in dark condition within genotype, n=17-39), Kruskal-Wallis, Dunn’s posthoc at hour 4, #p<0.05, ###p<0.001. Line graphs show mean and s.e.m. over time. *p<0.05, **p<0.01, ***p<0.001 (**a**,**d**,**e**,**g**,**i**). Scatter plot show mean +/-s.e.m. (**a**,**d**,**e**). Scale bar 50 μm (**c**,**f**,**h**). For full geno-types see Supplementary Table 1.

Postmated changes in female behavior are elicited by sex peptide (SP)^1,2,24^, a male-derived seminal fluid peptide and require female uterine primary sensory neurons (SPSNs)^13-15^ (Fig. 2b). To test the involvement of the SPSNs in postmated sucrose feeding changes, we genetically accessed SPSNs using three different genetic driver combinations (see methods). All genetic intersections consistently labeled at least 4 SPSNs (Fig. 2c, Extended Data Fig.1b,c) with no overlap of off-target neurons (Extended Data 1d), making them useful tools to investigate the role of the SPSNs in postmating feeding changes. We silenced the SPSNs in virgin females using constitutive silencers (either the potassium inward rectifier, KIR2.1, or the tetanus toxin light chain, TNT) to mimic the mated state and measured sucrose consumption. Virgin females with SPSNs silenced consumed significantly more sucrose than virgin controls (Fig. 2d, Extended data 1e,f), recapitulating the mated phenotype. This data argues that mated females increase sucrose consumption after mating via the SPSNs.

The canonical sex peptide pathway for postmated changes in female behavior is a three-layer circuit from SPSNs to SAGs to pC1^3,16,17^. As with the SPSNs, SAG and pC1 neurons are active in a virgin female and silenced in a mated female^3,16^. To precisely manipulate SAGs, we generated a new split-GAL4 line (SAG-SS3, Fig. 2m). Next, we acutely silenced SAGs by driving the expression of the green-light gated anion channelrhodopsin (GtACR1) with SAG-SS3. Monitoring consumption over time, we found that acutely silencing SAGs significantly increased sucrose consumption (Fig. 2n, Extended Data Fig. 2a). Although an off-target ascending neuron is also labeled in SAG-SS3, it is not responsible for changes in sucrose consumption (Extended Data Fig. 3), arguing that the increased sucrose consumption observed with SAG-SS3 is due to the activity of SAG neuron itself. Downstream of SAG, we found that acutely silencing pC1 also significantly increased sucrose consumption (Fig. 2o,p, Extended Data Fig. 2b-d), demonstrating that pC1 neurons regulate postmating sucrose feeding. Taken together, we conclude that the increased sugar consumption after mating is part of the repertoire of behavioral changes induced after copulation by sex peptide acting on the SPSN-SAG-pC1 mating status circuit.

**Fig. 3.**
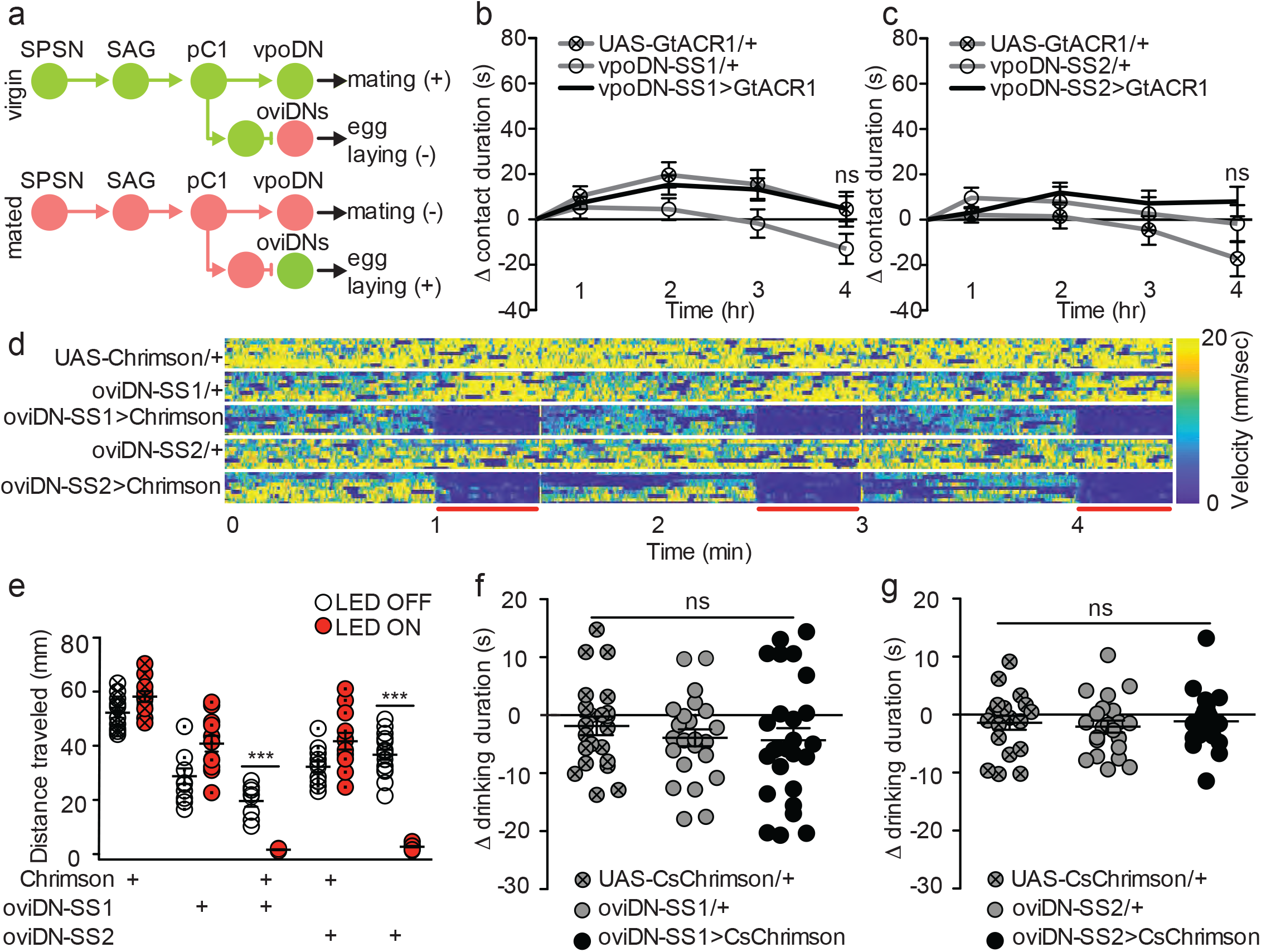
Descending neurons that regulate egg laying and sexual receptivity do not influence sucrose consumption. **a**, Schematic of mating circuit with neural outputs of pC1 and resulting postmating response. Green circles indicate active neurons, salmon circles indicate silenced neurons. **b**-**c**, Difference in drinking time of indicated genotype in FLIC over 4 hrs. Each data point represents normalized drinking time of a single female in light condition (drinking time minus the daily average drinking time of all females in dark condition within genotype, n=20-24), Kruskal-Wallis. Line graph shows mean and s.e.m. over time. **d**, Velocity heatmap upon transient activation in individual walking flies. Red bars indicate light ON (n=8-16). **e**, Total distance traveled in locomotor assay (**d**). ON condition (red) includes 3 30-sec light ON conditions. OFF condition (white) includes 3-30sec light OFF conditions immediately preceding each ON condition, Mann Whitney, ***p<0.001. Aligned dot plot show mean +/-s.e.m. **f**-**g**, Drinking time of females exposed to light normalized to dark controls of indicated genotype in FLIC over 4-hrs (n=19-26), 1way ANOVA, Bonferroni’s posthoc. ns = not significant. Scatter plot show mean +/-s.e.m. (**e**-**g**), For full genotypes see Supplementary Data Table 1.

Postsynaptic to pC1, the mating status circuit splits into separate neural pathways that mediate different behavioral subprograms, including circuits that elicit changes in sexual receptivity^17^ or egg-laying^16^. We hypothesized that the neural circuit supporting the postmated increase in sucrose consumption could either follow one of the established circuits or diverge after pC1. To test if sucrose feeding is executed by previously identified postmating subcircuits, we optogenetically manipulated the activity of the descending neurons (DNs) for sexual receptivity (vpoDNs) or egg laying (oviDNs) to mimic the mated state (Fig. 3a) and measured sucrose consumption. We found no difference in sucrose consumption upon manipulation of vpoDNs or oviDNs (Fig. 3b,c,f,g, Extended Data Fig. 2e-h). We note that we measured sucrose intake in immobilized flies for oviDN manipulations, as activating oviDNs caused females to stop walking (Fig. 3d,e), likely a natural component of egg laying. These studies argue that neither of the previously identified circuits responsible for a single postmating response, egg laying and sexual receptivity, mediate postmating sucrose feeding.

To identify novel pathways that mediate changes in mated feeding behavior, we examined other major outputs of the mating status circuit. We identified postsynaptic partners of SAG and pC1 using automated analysis^25,26^ of a fly brain electron microscopy (EM) connectome^27,28^. This approach identified three pCd-2 neurons^29,30^, pCd-2a, pCd-2b, and pCd-2c (Fig. 4a, Extended Data Fig. 4a), that are strongly connected to SAG (39 synapses) and pC1 (158 synapses). These neurons are promising candidates to mediate postmating feeding changes because their processes descend into the prow region of the Subesophageal Zone (SEZ, Fig. 4b,c), a brain region implicated in energy homeostasis and feeding^31^. As pC1 and SAG are cholinergic^3,16^, pCd-2 cells likely receive excitatory signals from these two upstream synaptic partners and therefore would be predicted to be less active in mated females (Fig. 4d). We generated split-GAL4 lines to genetically access pCd-2a and pCd-2b (Fig. 4e,f) and found that both neurons are female-specific (Extended Data Fig. 4b), consistent with a role in female postmating behavior. Optogenetically silencing either pCd-2a or pCd-2b significantly increased sucrose consumption, recapitulating the mated state (Fig. 4h,I, Extended Data Fig. 5c-j). Importantly, silencing pCd-2a or pCd-2b did not influence egg laying (Fig. 4g, Extended Data Fig. 5a,b), demonstrating separation of circuits for increased sucrose consumption from other behaviors downstream of pC1.

**Fig. 4.**
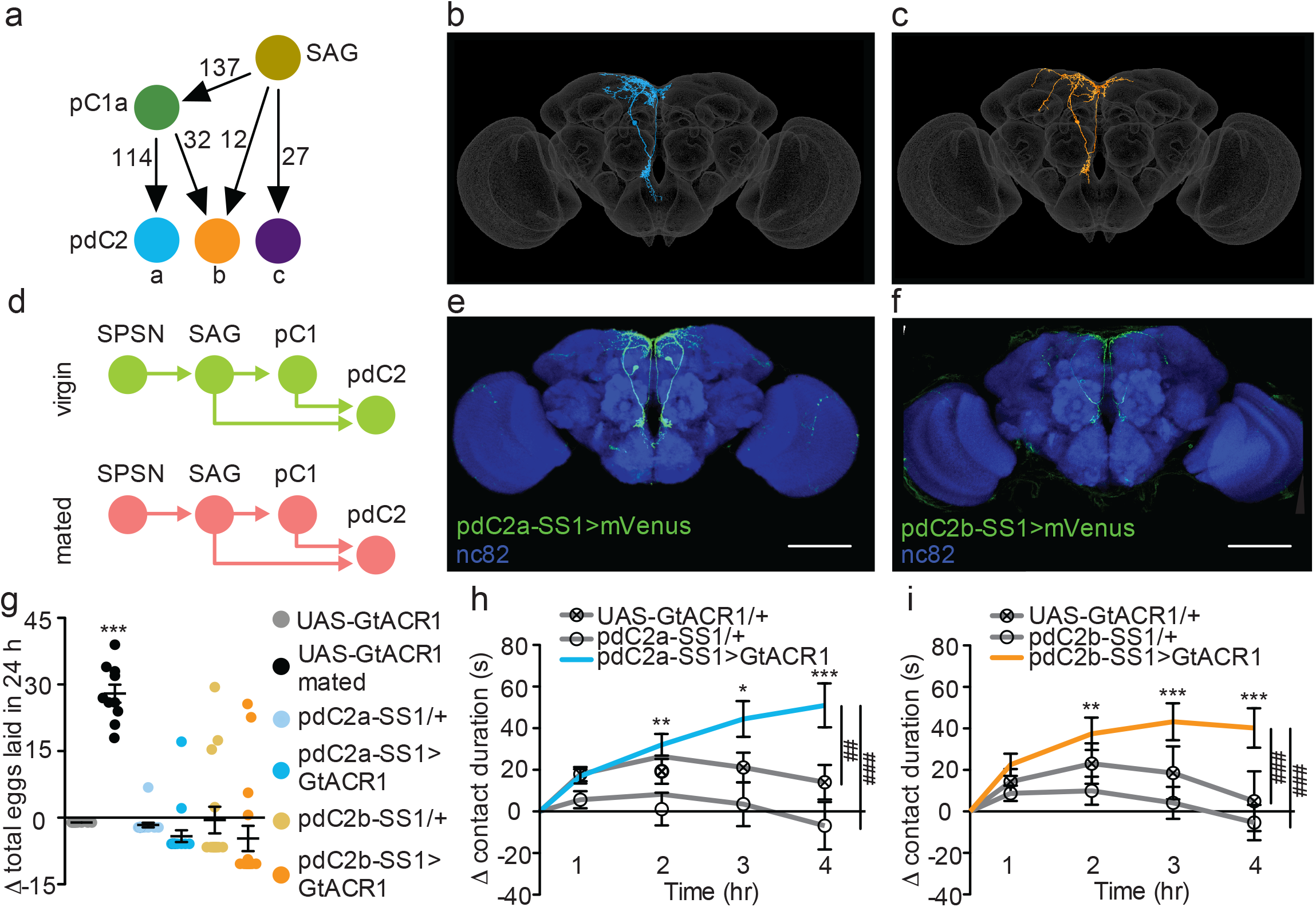
pCd-2 neural outputs of SAG and pC1 mediate postmating sucrose consumption. **a**, Schematic of neural connectivity of the mating status circuit and pCd-2 cells. Arrows represent direct connection and number indicates number of synapses. **b**-**c**, Electron microscopy reconstruction of pCd-2a,b neurons. **d**, Model of circuit activity in virgin and mated state. Green circles indicate active neurons, salmon circles indicate silenced neurons. **e**-**f**, Confocal images of pCd-2a-SS1 and pCd-2b-SS1 brains, stained to reveal pCd-2 projection pattern (UAS-GFP, green) and all synapses (nc82, blue). Scale bar 50 μm. **g**, Difference in eggs laid in 24-hr of females exposed to light normalized to dark controls (n=10-20). For “UAS-GtACR1 mated” group, data set represents eggs laid by mated females normalized to virgin females of the same genotype. Scatter plot show mean +/-s.e.m., one sample t-test. **h**-**i**, Drinking time of females exposed to light normalized to dark controls of indicated genotype in FLIC over 4-hrs (n=22-26). Line graph show mean and s.e.m. over time, Kruskal-Wallis, Dunn’s posthoc at hour 4, ##p<0.01, ###p<0.001. *p<0.05, **p<0.01, ***p<0.001. For full genotypes see Supplementary Table 1.

**Fig. 5.**
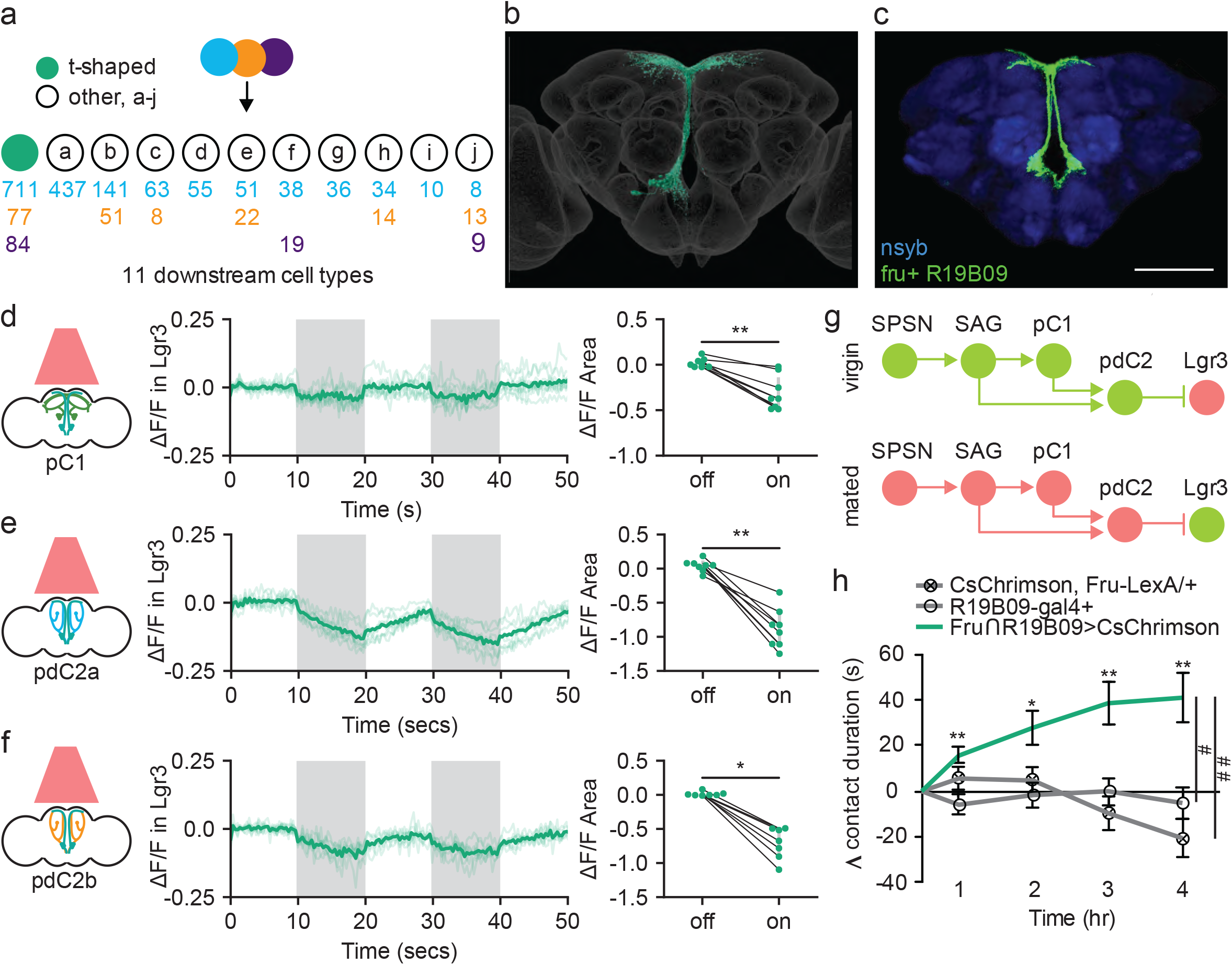
Sexually dimorphic neuroendocrine Lgr3+ cells receive mating status signals via pCd-2 neurons and regulate sucrose consumption in mated females. **a**, Downstream neuronal cell types of pCd-2 cells identified by electron microscopy reconstruction, showing the number of pCd-2 output synapses. T-shape cell type represented with turquoise circles, other cell types represented with white circles (a-j). **b**, Electron microscopy reconstruction of t-shape neurons. **c**, Confocal image of fru+ Lgr3-expressing cells in medial brain (UAS-CD8:GFP, green) and all synapses (nc82, blue). Scale bar 50 μm. **d**-**f**, Calcium responses of Lgr3-expressing cell bodies in the prow region of the brain to activation of upstream pC1 (**d**), pCd-2a (**e**), or pCd-2b (**f**) neurons (n=7-8) and the analysis of area under the curve, Wilcoxon Rank Test, scatter plot show mean +/-s.e.m. (grey bar). **g**, Neural model for the coordination of mating status and sucrose intake. Green circles indicate active neurons, salmon circles indicate silenced neurons. **h**, Drinking time of females exposed to light normalized to dark controls of indicated genotype in FLIC over 4-hrs (n=10-15). Line graph shows mean and s.e.m. over time, Kruskal-Wallis, Dunn’s posthoc at hour 4, #p<0.05, ##p<0.01. *p<0.05, **p<0.01, ***p<0.001. For full genotypes see Supplementary Table 1.

To examine how pC1 influences sucrose consumption, we used EM analyses and identified the major downstream targets of pCd-2 as t-shaped cells (711 synapses) having processes in the SEZ prow region and a neuroendocrine center, the pars intercerebralis, with terminal arbors similar to insulin producing cells and other median secretory cells^32^ (Fig. 5a, Extended Data Fig. 6). Based on anatomical similarity, these cells are Lgr3+ (leucine-rich repeat G-protein-coupled receptor^33^) medial bundle neurons (mBNs) (Fig. 5b,c). Lgr3 expression is sexually dimorphic, found specifically in the central brain and abdominal ganglia of females^34^, consistent with a role in female behavior. Moreover, Lgr3 is a receptor for the *Drosophila* insulin-like peptide 8 (Dilp8^35-37^), and both Lgr3 and Dilp8 are known to regulate feeding^38^. These findings suggest that the Lgr3+ mBNs may represent the site where mating status circuits interact with hunger/satiety circuits to regulate feeding.

To test this hypothesis, we first examined whether Lgr3+ mBNs are functionally connected to pC1, pCd-2a, or pCd-2b by optogenetically activating these populations while simultaneously measuring calcium responses in Lgr3+ neurons. In all cases, neural activation caused significant calcium decreases in Lgr3+ neurons (Fig. 5d-f), validating Lgr3+ mBNs as targets of the mating status pathway. As pC1 is inhibited in the mated state, our studies argue that Lgr3+ mBNs are more active in mated females (Fig. 5g). To examine whether activity in Lgr3+ mBNs influences consumption, we targeted this population with a genetic intersectional approach (R19B09∩fru^34^, Fig. 5c), optogenetically activated Lgr3+ mBNs, and monitored sucrose consumption. We found that increased activity in Lgr3+ mBNs promotes sucrose consumption, mimicking the mated state (Fig. 5h). Together, these studies reveal that Lgr3+ mBNs receive inputs from the canonical postmating pathway and are part of a neuroendocrine pathway that regulates sucrose consumption in postmated females.

In conclusion, we report a new postmated response of increased sucrose consumption, which independent of caloric deficiencies caused by egg production and anticipatory in nature. Our findings provide a neural mechanism for the coordination of mating state and sucrose consumption (Fig. 5g). In virgins, increased activity of SAG and pC1 potentiates pCd-2-mediated inhibition of Lgr3+ cells within the endocrine center of the brain, resulting in low sugar intake. After mating, the activity of SAG and pC1 is reduced^3,16^, decreasing the activity of pCd-2 and alleviating the inhibition from *Lgr3+* cells, increasing sucrose intake.

The sexually dimorphic Lgr3+ neurons are the nexus integrating mating status and neuroendocrine hunger signaling. Lgr3 is a leucine-rich repeat containing G-protein coupled receptor, which binds Dilp8 to regulate feeding in flies. Expression levels of Lgr3 and Dilp8 rise in fed flies and mutations in either gene increase feeding, arguing that Dilp8 and Lgr3 are satiety factors^38^. Our calcium imaging and behavioral studies of Lgr3+ mBNs indicate that they are more active in a mated state and promote sucrose consumption. Together, these data suggest a model for how Lgr3+ mBN neurons integrate signals from the mating status and hunger/satiety systems. In virgin or satiated female flies, circulating DILP8 and pCd-2 activity, respectively, inhibit Lgr3+ mBNs (Fig. 5g), reducing sugar feeding. Conversely, in mated or hungry flies, Lgr3+ mBNs lack the inhibitory signal from pCd-2 neurons or circulating Dilp8, respectively, resulting in increased sugar consumption. Hence, the mated state may be viewed mechanistically as an additional “hunger” signal within this system.

Thus, we report that an elevated appetite for sucrose is an important behavioral subprogram elicited by mating and executed by female-specific circuitry, shifting the physiology of a mated female. The activation of hunger centers by the mating status circuit provides a neural mechanism that anticipates the large energetic demands associated with offspring production and increases caloric intake, promoting reproductive success.

---

**Extended Data Fig. 1.**
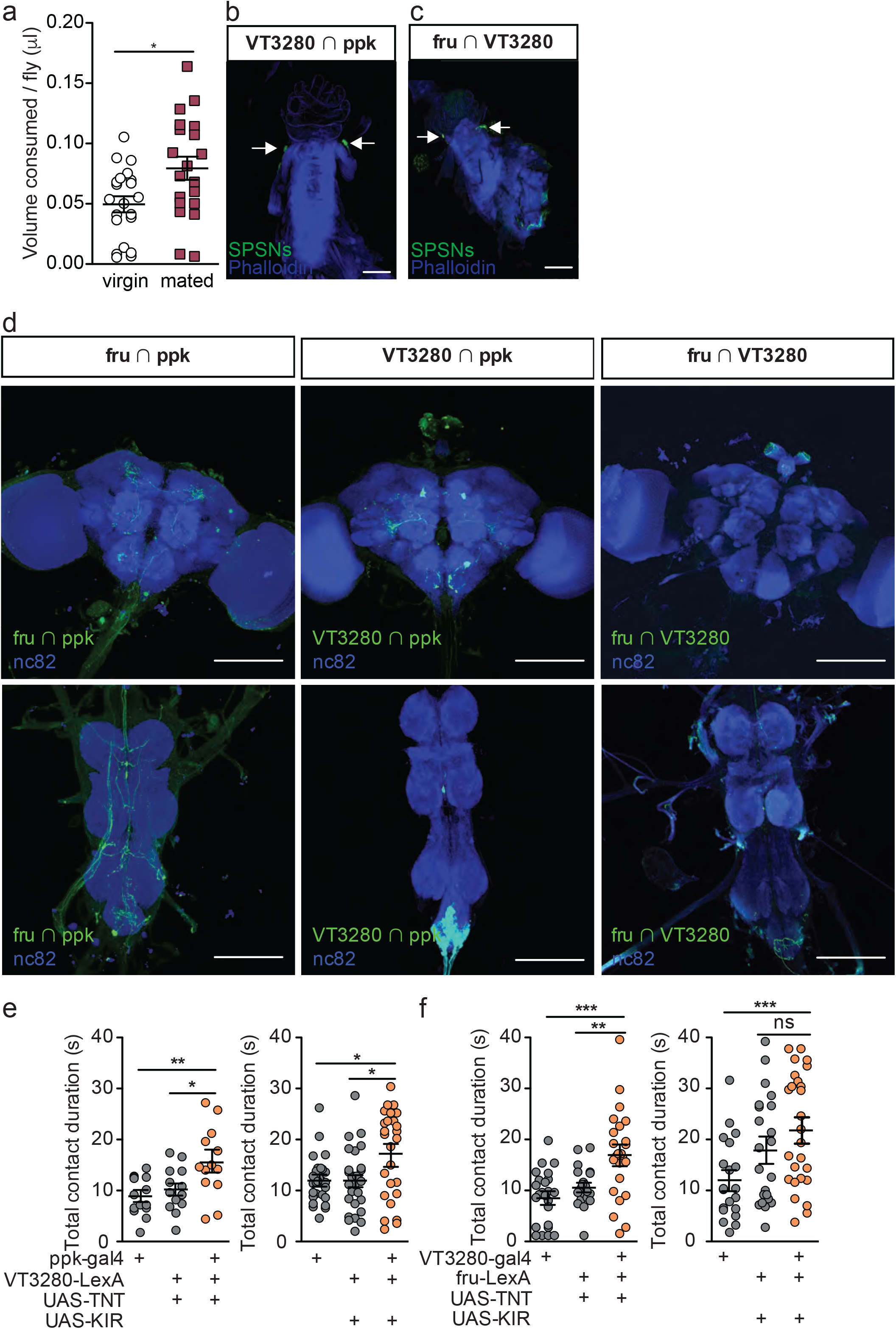
Three independent SPSN lines regulate postmated sucrose consumption. **a**, Total volume of 50mM sucrose consumed per fly in CAFE assay by eggless virgin (n=21) and eggless mated females (n=20). *p<0.05, unpaired t-test. **b**-**c**, Confocal image of uterus stained to reveal SPSNs (UAS-CD8:GFP, green, arrowhead indicating cell bodies) and muscle tissue (phalloidin, blue). **d**, Confocal images of brain (top) and ventral nerve cord (bottom) of genetic intersections accessing SPSNs stained to reveal off-target neurons in the central nervous system (UAS-GFP; green) and all synapses (nc82; blue). **e**-**f**, Total drinking time in FLIC in 20-min assay of virgin females of indicated genotype (n=19-41). Kruskal-Wallis with Dunn’s Multiple Comparison Test revealed differences between silenced females and controls. Scatter plot show mean +/-s.e.m. (**a**,**e**-**f**). Scale bar 50 μm (**b**-**d**).

**Extended Data Fig. 2.**
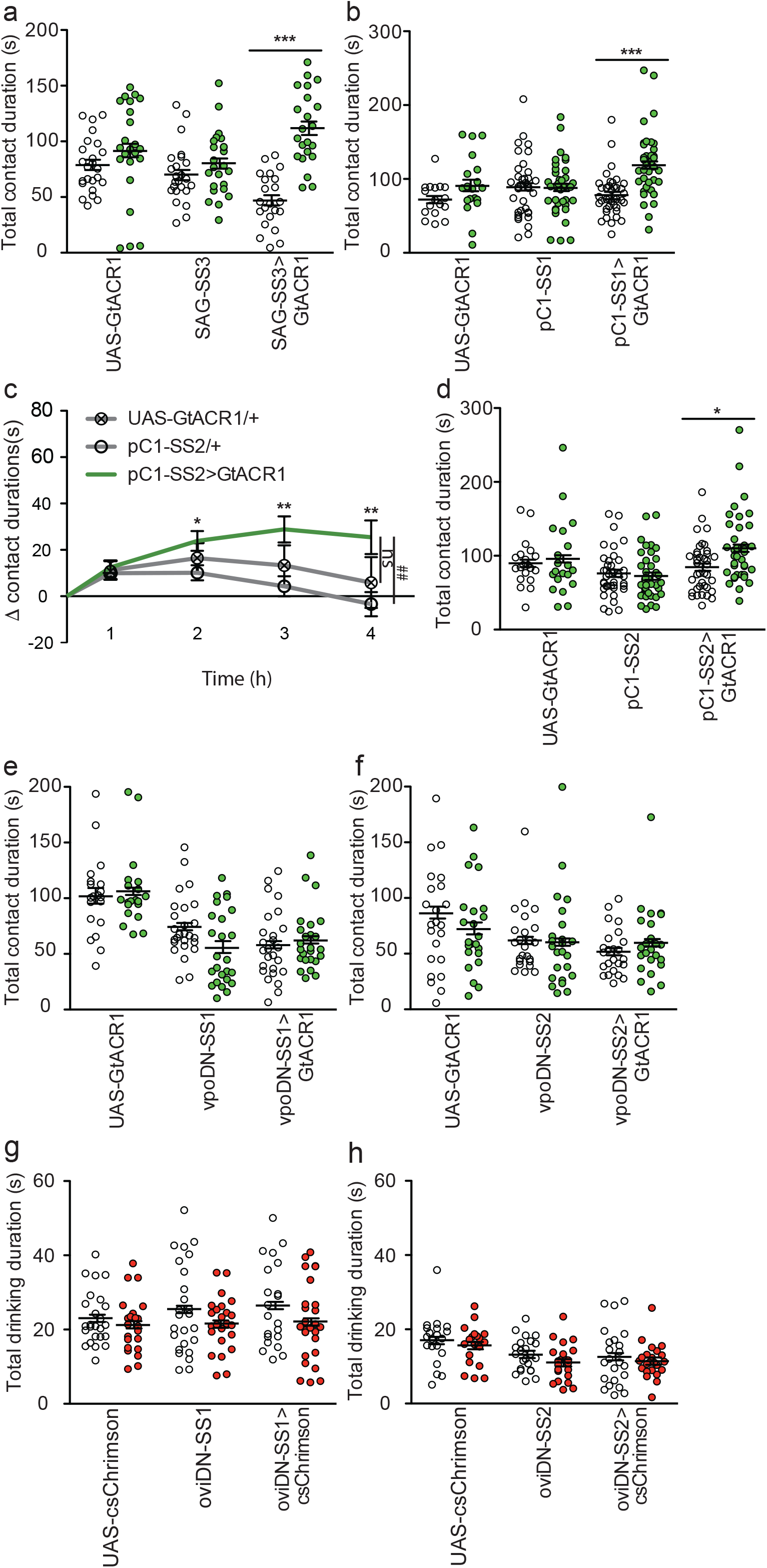
Contact duration times of virgin and mated females upon optogenetic manipulation of neurons in the mating status circuit. **a-b**, Total drinking time in FLIC in 4-hr assay of females of indicated genotype and light condition. White circles are dark condition, green circles are green light condition, n=20-32. Data was analyzed with 2way ANOVA and Bonferroni posttests, ***p<0.001. **c**, Drinking time of females exposed to light normalized to dark controls of indicated genotype in FLIC over 4-hrs, n=20-24. Line graphs show mean and s.e.m. over time, Kruskal-Wallis, Dunn’s posthoc at hour 4, ##p<0.01, ns = not significant. **d-f**, Total drinking time in FLIC in 4-hr assay of females of indicated genotype and light condition. White circles are dark condition, green circles are green light condition, n=20-32, 2way ANOVA and Bonferroni posttests. **g-h**, Total drinking time in TCA of females of indicated genotype and light condition. White circles are dark condition, red circles are red light condition, n=20-32, 2way ANOVA and Bonferroni posttests, *p<0.05, **p<0.01. Scatter plot show mean +/-s.e.m. (**a**-**b**,**d**-**h**).

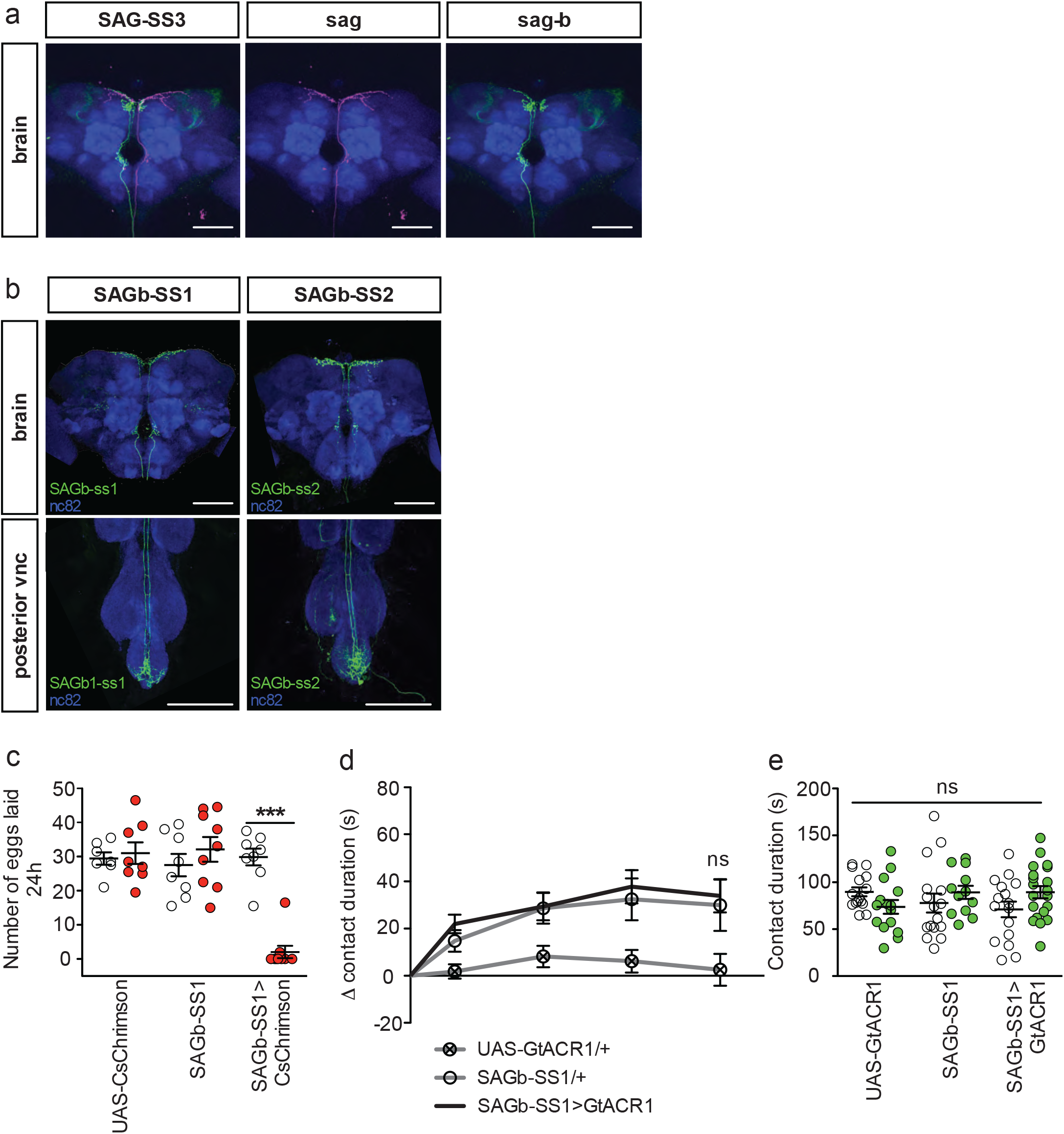

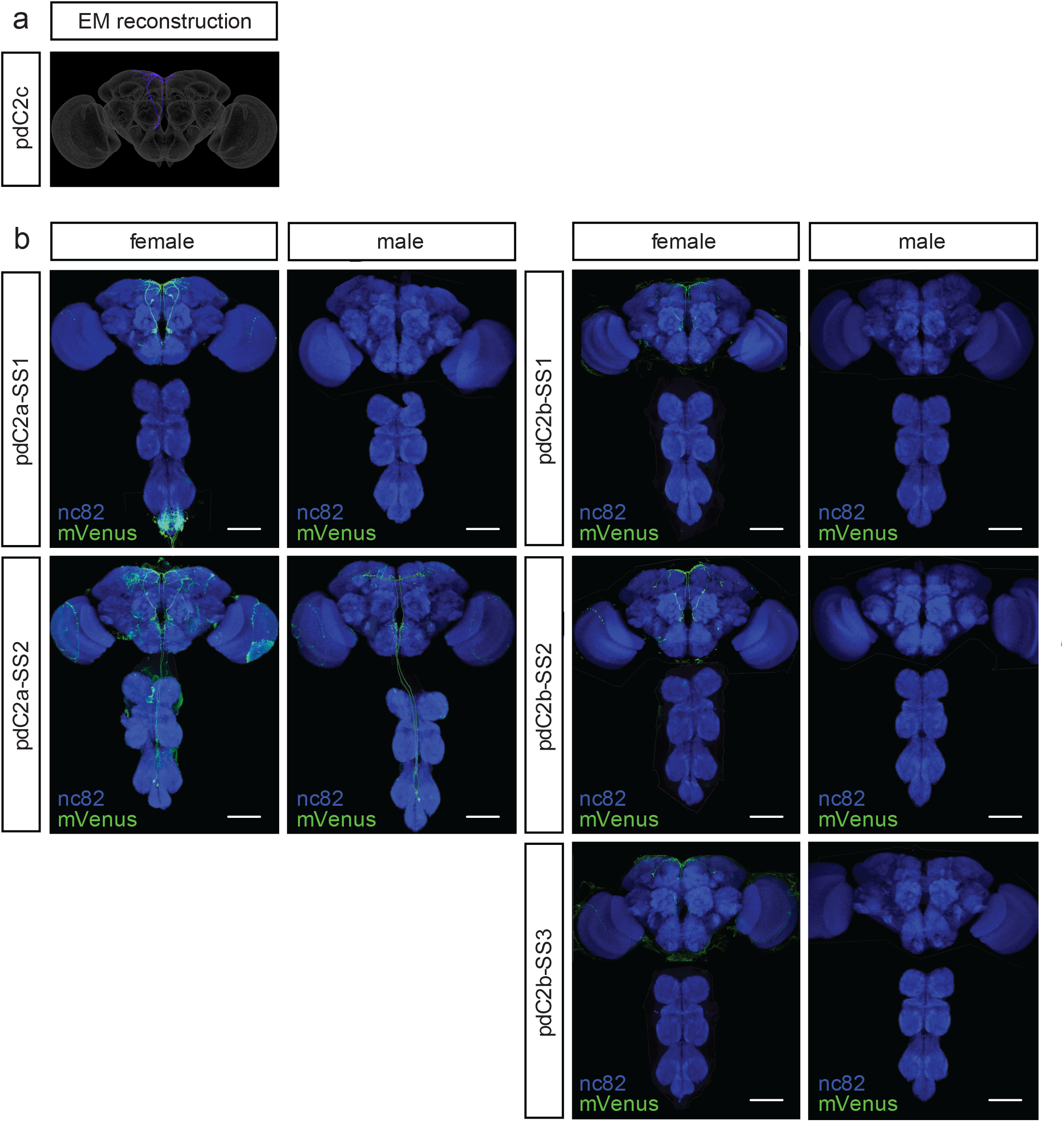

**Extended Data Fig. 5.**
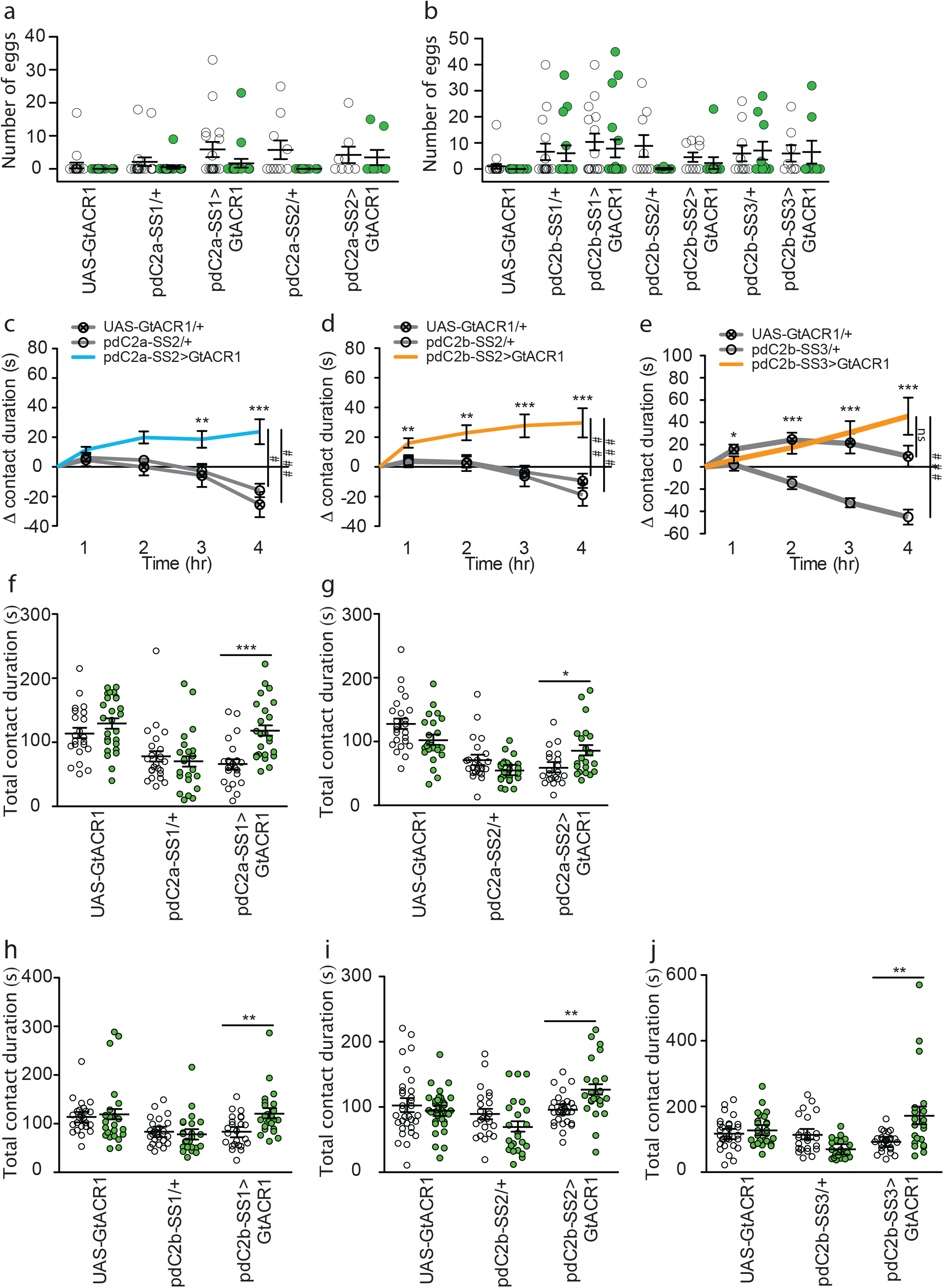
Independent pCd-2a and pCd-2b lines regulate postmated sucrose consumption. **a**-**b**, Number of eggs laid in 24 hours by virgin females of indicated genotype while exposed to green light (green) or no light (white). 2way ANOVA analysis revealed no significant differences between groups (n=9-20). **c**-**e**, Drinking time of females exposed to light normalized to dark controls of indicated genotype in FLIC over 4-hrs (n=19-36), Kruskal-Wallis, Dunn’s posthoc at hour 4, ###p<0.001. Line graph shows mean and s.e.m. over time. **f**-**j** Total drinking time in FLIC in 4-hr assay (n=20-24). Analyzed with a 2way ANOVA and Bonferroni posttests (n=19-36). *p<0.05, **p<0.01, ***p<0.001, ns = not significant. Scatter plot show mean +/-s.e.m. (**a**-**b**,**f**-**j**).

**Extended Data Fig. 6.**
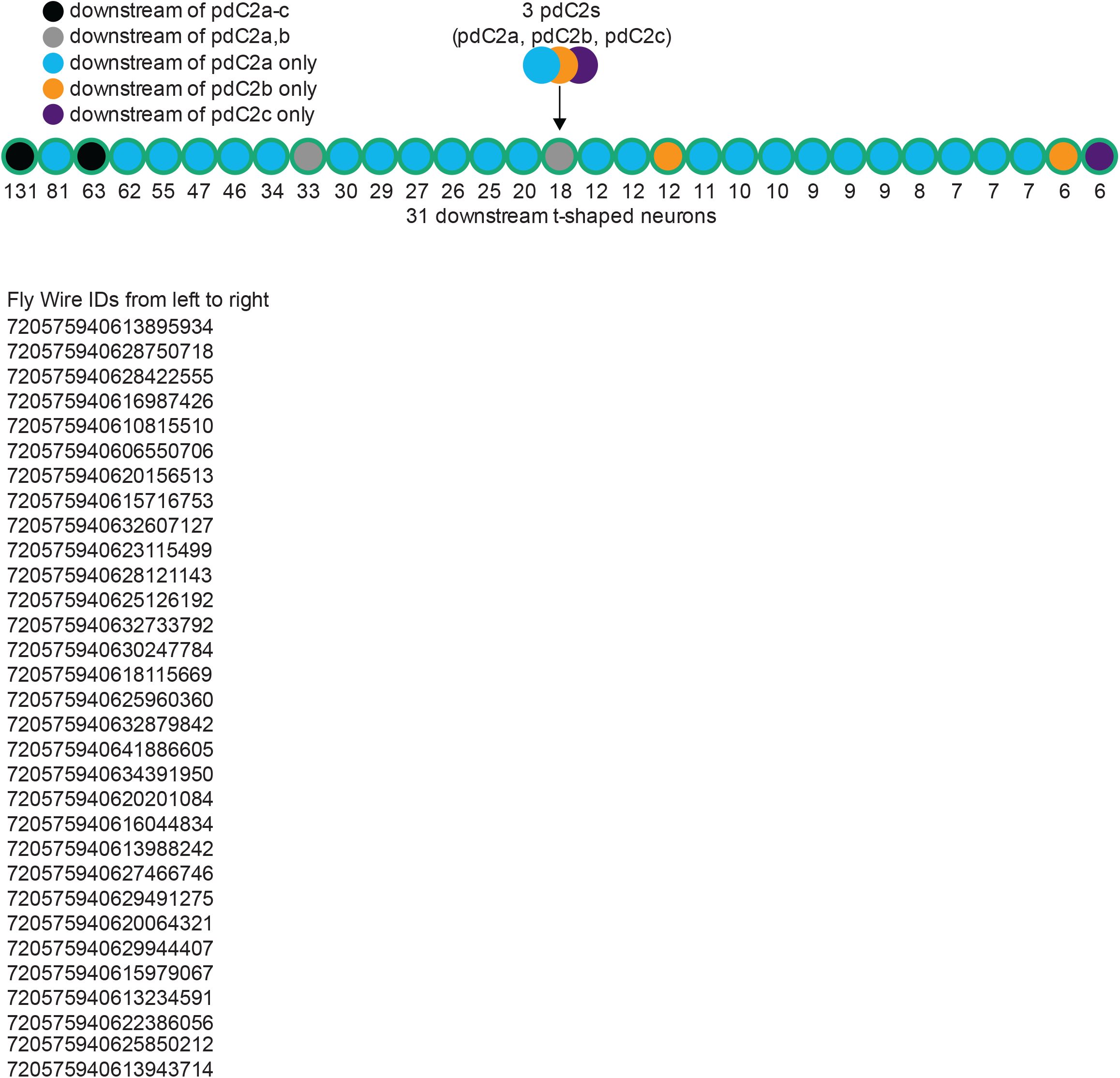
Post-synapatic targets of pCd-2. Schematic of neural connectivity of pCd-2 cells and downstream t-shape neurons identified by electron microscopy reconstruction. Number indicates number of synapses.

## Methods

### Rearing conditions and strains

Flies were reared on standard cornmeal-agar-molasses medium, at 25°C, 65% humidity on a 12-hr-light-dark cycle. Flies used in optogenetic assays were reared on food containing 0.25mM all-trans-retinal (Sigma-Aldrich) in darkness, before and after eclosion.

Flies were collected under CO_2_ 1-8h after eclosion and housed in same-sex groups. To generate mated females, 15 virgin females and 5 males were paired 3-5 days post-eclosion and group housed for 72 hours. To generate recently mated females, a single virgin female was paired with a single male for 2 hours. If mating was observed, the recently mated female was then immediately placed into indicated assay. To generate 24-hr mated females, a single virgin female was paired with a single male for 2 hours. If mating was observed, the recently mated female was then group housed with other recently mated females until testing. To generate egg-less females we used either ovoD mutant females^23^ or hsp70-bam females^22^ following the heat-shock protocol^7^.

### Capillary feeding (CAFE) assay

Capillary feeding arenas were modified from the original^20^. Briefly, we equipped a plastic vial, containing a wet kimwipe, with an altered rubber stopper lid housing two truncated 200-ml pipette tips whereby the capillaries can be inserted through. The capillaries (calibrated glass micropipettes, 5 ml) were loaded with 50mM sucrose solution containing 0.25 mg/ml blue dye. To measure sucrose consumption, 5 virgin or mated females were gently aspirated into an arena for 24 hours, the displacement of the meniscus was measured, and the volume per fly was calculated (1mm = 0.038ml).

### Fly liquid interaction counter (FLIC)

Females of indicated genotype or mating status were first wet-starved for 12-18 hours. Following this, females were then gently aspirated into the fly liquid food interaction counter (FLIC^18^), which was pre-filled with sucrose solution of indicated concentration. The voltage from each well was continuously recorded and a .csv file was produced. Significant changes in the voltage indicated that the fly was contacting the liquid.

Contact duration acted as a proxy for consumption time. Based on a video analysis of flies drinking in FLIC in our lab, we modified the parameters in R to better model consumption. The new parameters are as follows: Feeding.Interval.Minimum = 10, Feeding.Threshold.Value = 10, Tasting.Threshold.Interval = c(01,02), Feeding.Minevents = 6, Signal.Threshold = 10.

For optogenetic control of neural activity prior to FLIC, we placed vials containing flies into a custom made light box (20cm x 10cm x 30cm) equipped with a string of 150 LED lights programmed with an Arduino to have green light constantly on.

For optogenetic control of neural activity while flies were in the FLIC, we designed and constructed a 12-chamber lid equipped with either 3 red LEDs (617nm, Luxeon Star, catalog number SP-01-E6) or 3 green LEDs (530nm, Luxeon Star, catalog number SP-01-G4). The 12-chamber cover was made from half of a 24-well plate, which was trimmed to fit. Lights were attached onto half of the 24-well plate lid and fitted atop the chamber cover.

### Proboscis extension response (PER) assay

Females of indicated mating status were either wet-starved for 12-18 hours (starved) or kept on standard food (fed). Flies were then anesthetized using CO_2_ and fixed to a glass slide with No More Nails polish and then allowed to recover for 2 hours in a humidified box. Immediately before testing, flies were given water until they no longer responded to 5 consecutive presentations. Flies were then presented 3 times with sucrose of an indicated concentration and proboscis extension was recorded.

### Temporal consumption assay (TCA)

Flies were starved for 18 hours in darkness, anesthetized using CO_2_ and then fixed to a glass slide with No More Nails polish with limited light and allowed to recover for 2 hours in a humidified red light LED box or a humidified dark box.

Immediately before testing, flies were given water until they no longer responded to 5 consecutive presentations. In testing, flies were presented with 250 mM sucrose and consumption time was recorded. Each trial ended when the fly was presented with the tastant and no longer responded to 5 consecutive presentations. Flies in the dark condition were given water and tested in low light conditions. Flies in red light condition were given water and tested in the presence of red light.

### Egg laying assay

7-day old females were placed in a vial with standard food and placed into a custom-made light box (20cm x 10cm x 30cm) that was equipped with a string of 150 LED lights programmed with an Arduino. In neural activation tests, red light was transiently on (1 second on/1 second off). In neural silencing tests, green light was constantly on. Half of the females of each genotype were placed in vials wrapped in foil to function as no-light condition controls. After 24 hours, females were removed and eggs were immediately counted.

### Locomotor assay

Locomotor assays were conducted as described^39^.

### In vivo calcium imaging with taste stimulation

Protocol was adapted^31^. Virgin and mated females aged 14-20 days and mated females were co-housed with Canton-S males. Females were food-deprived for 18-24 hours prior to imaging. A window was made in the head of the fly to allow for visualization of the SEZ and a drop of ∼250 mOsmo AHL solution (https://www.sciencedirect.com/science/article/pii/S0092867403000047) was added to the head and imaging was immediately performed on a fixed-stage 3i spinning disk confocal microscope with a piezo drive and a 20x water objective (1.6x optical zoom). A 250 mM sucrose solution was delivered to the proboscis using a glass capillary. To analyze these images, a maximum intensity projection encompassing the arbors of Gr64f neurons was made using Fiji. Large ROIs were drawn manually corresponding to the GCaMP expression and a large ROI was drawn in an adjacent region to measure background autofluorescence. Mean fluorescence levels from the background ROI were subtracted from the Gr64f ROIs at each time point, resulting in the fluorescence trace over time: F(t). delta F/F was measured as follows: (F(t) – F(0)) / F(0), where F(0) was the average of 10 time points before stimulation with sucrose and F(t) was the average background subtracted increases in fluorescence during sucrose stimulation. Area under the curve for 10 frames before stimulation and during stimulation was approximated with the trapezoidal rule in Python using the NumPy.trapz function. Statistical analysis was carried out in Python using the SciPy package, version 1.7.3 (https://www.nature.com/articles/s41592-019-0686-2). deltaF/F0 images were created as described^40^.

### In vivo functional connectivity experiments

In vivo sample preparation for calcium imaging was performed as described^31^. Prior to imaging, female flies were aged for 2 weeks and then food-deprived in a vial containing a distilled water-soaked kimwipe for 18–24 hours prior to imaging. Female flies were housed with Canton-S males throughout aging and food-deprivation.

Following dissection, samples were bathed in ∼250 mOsmo AHL solution (https://www.sciencedirect.com/science/article/pii/S0092867403000047) and imaged immediately using a 2-photon microscope. Volumetric images of the prow region were acquired with 920 nm light at 2.9 Hz with resonant scanning and a piezo-driven objective. During imaging, a custom ScanImage plugin was used to deliver two 10-second pulses of 660 nm light through the objective at a 10-second interval to excite CsChrimson.

Image and statistical analysis of functional connectivity experiments was performed using Fiji (https://www.nature.com/articles/nmeth.2019), CircuitCatcher (a customized python program by Daniel Bushey), and Python (Python Software Foundation, https://www.python.org/). Using Fiji, image stacks for each time point were first maximum intensity projected. Then, using CircuitCatcher, an ROI containing Lrg3+ mBN cell bodies and a background ROI of brain tissue were selected and the average fluorescence intensity for each ROI at each timepoint was retrieved. Then, in Python, background subtraction was carried out for each timepoint (F_t_) and initial fluorescence intensity (F_initial_) was calculated as the mean corrected average fluorescence intensity for frames 1-15. Then, ΔF/F was calculated using the following formula: F_t_-F_initial_/F_initial_. Area under the curve for before (off; frames 1-29) and during (on; frames 30-58) light stimulation was approximated with the trapezoidal rule in Python using the NumPy.trapz function. Statistical analysis of functional connectivity experiments was carried out in Python using the SciPy package, version 1.7.3 (https://www.nature.com/articles/s41592-019-0686-2).

### Dissections and Immunohistochemistry

All CNS dissections and immunostaining (unless directly addressed) were performed following the detailed instructions found at https://www.janelia.org/project-team/flylight/protocols. To image split-GAL4 lines and intersections, we used the following primary antibodies: mouse α-Brp (nc82, DSHB, University of Iowa, USA) at 1:40, chicken α-GFP (Invitrogen, A10262) at 1:1000; and the following secondary antibodies: goat α-mouse AF647 (Invitrogen, A21236) at 1:500, goat α-chicken AF488 (Life Technologies, A11039) at 1:500. Multi-Color Flip-Out fly generation was performed following the protocol^41^. For imaging, we used the following primary antibodies: mouse α-Brp (nc82, DSHB, Univerisity of Iowa, USA) at 1:40, rabbit α-HA (rabbit α-HA (Cell Signaling Technology, C29F4) at 1:1000, rat α-Flag (Novus Biologicals, NBP1-06712) at 1:1000; and the following secondary antibodies: goat α-mouse AF647 (Invitrogen, A21236) at 1:500, goat α-rabbit AF488 (Invitrogen, A11034) at 1:500, and goat α-rat AF568 (Invitrogen, A11077) at 1:500.

Reproductive tract dissections and staining was performed as described^42^. As the primary antibody, we used chicken α-GFP (Invitrogen, A10262) at 1:1000 followed by goat α-chicken AF488 (Life Technologies, A11039) at 1:500 with the addition of Alexa Fluor 633 phalloidin (ThermoFisher Scientific) to visualize the muscle-lined reproductive tract (1:500).

### Confocal imaging

Samples were imaged on a LSM710 confocal microscope (Zeiss) with a Plan-Apochromate 20×0.8 M27 objective and images were prepared in Fiji.

### Electron Microscopy neural reconstructions and connectivity

SAG was previously partially reconstructed^16^ in the Full Adult Female Brain (FAFB^27^) electron microscopy dataset using the CATMAID software^43^. We completed additional tracing of this neuron by continuing branches with a combination of both manual and assisted tracing. Manual tracing consisted of generating a skeleton of the neuron by following the neuron and marking the center of each branch. Assisted tracing consisted of joining and proofreading pre-assembled skeleton fragments with automated segmentation^44^. In addition to the skeleton tracing, new chemical synapses were also annotated as previously described^27^. Finally, downstream synaptic targets of SAG were then traced out from these additional locations using both manual and assisted tracing techniques as described above.

Neurons traced in CATMAID, including SAG, pC1, egg-laying circuit neurons^16^ and sexual receptivity circuit neurons^17^, were all located in Flywire (flywire.ai), which uses the same EM electron microscopy dataset^27^. To identify synaptic partners, we used connectome annotation versioning engine (CAVE^25,26^) using a cleft score cutoff of 100 to generate synapses of relatively high confidence^26^.

### Intersectional method to access SPSNs

Based on previous reports, three genetic drivers label the SPSN (VT003280[3], fru[14,15] and ppk[14,15]) with driver lines available in two separate binary expression systems (UAS-GAL4 and LexA-LexAop). With the use of a conditional reporter line (UAS < stop < GFP) and in combination with inducible FLP line (LexAopFLP) three combinations were produced: fru-LexA∩ppk-GAL4, fru-LexA∩VT3280-GAL4, and VT3280-LexA∩ppk-GAL4. Images of the female reproductive tract were used to confirm labeling of the SPSNs and images of the brain to determine off-target neural expression. With the use of a conditional effector line (UAS < stop < KIR or UAS < stop < TNT), SPSNs were silenced and females (along with genetic controls) were tested for changes in sucrose consumption.

### Split-GAL4 screening and stabilization

#### pCd-2 split generation

Using the FlyEM Hemibrain V1.2.1 dataset via NeuPRINT, we queried “SAG” and explored the output neuron list, identifying three pCd-2 neurons per hemisphere (SMP286, SMP287, SMP288). From here, we used Neuron Bridge to manually explore the light microscopy matches and generated a list of possible hemi-driver that labeled this cell type. The expression pattern of the p65ADZp and ZpGAL4DBD combinations were examined by driving the expression of UAS reporter (20xUAS-CsChrimson-mVenus in attP18). Combinations with specific expression in pCd-2 neurons were stabilized.

#### SAGb split generation

Using Neuron Bridge, we queried “VT050405” (the genetic construct used as the p65ADZp hemidriver in SAG-SS3) and identified multiple “Body Ids” that morphologically matched the non-SAG ascending cell type. From here we generated a “Color Depth Search” to create a list of possible hemi-driver that labeled this cell type. The expression pattern of the p65ADZp and ZpGAL4DBD combinations were examined by driving the expression of UAS reporter (20xUAS-CsChrimson-mVenus in attP18) and combinations with specific expression in SAGb neurons were stabilized.

### Statistical analysis

Statistical tests for all experiments, with the exception of in vivo calcium imaging and in vivo functional connectivity experiments, were performed in GraphPad Prism. For two- and three-group comparisons, data was first tested for normality with the KS normality test (alpha = 0.05). If all groups passed then groups would be compared with a parametric test (t-test or One-Way ANOVA, respectively). If at least one group did not pass, groups were compared with a non-parametric version (Mann-Whitney test or Kruskal-Wallis test, respectively). For all multi-factorial designs, a Two-Way ANOVA was performed with a Bonferroni post-hoc test. All statistical tests, significance levels, and number of data points (N) are specified in the figure legend.

### Normalized data sets

Most datasets from optogenetic assays were normalized within each genotype. To generate this normalized dataset, data from females within the no light condition was averaged, creating a “no-light mean” for each genotype. This value was subtracted from each individual female within the light condition of the corresponding genotype. This dataset was then graphed and statistical analysis was performed as outlined above.

---

**Supplementary Table 1.**
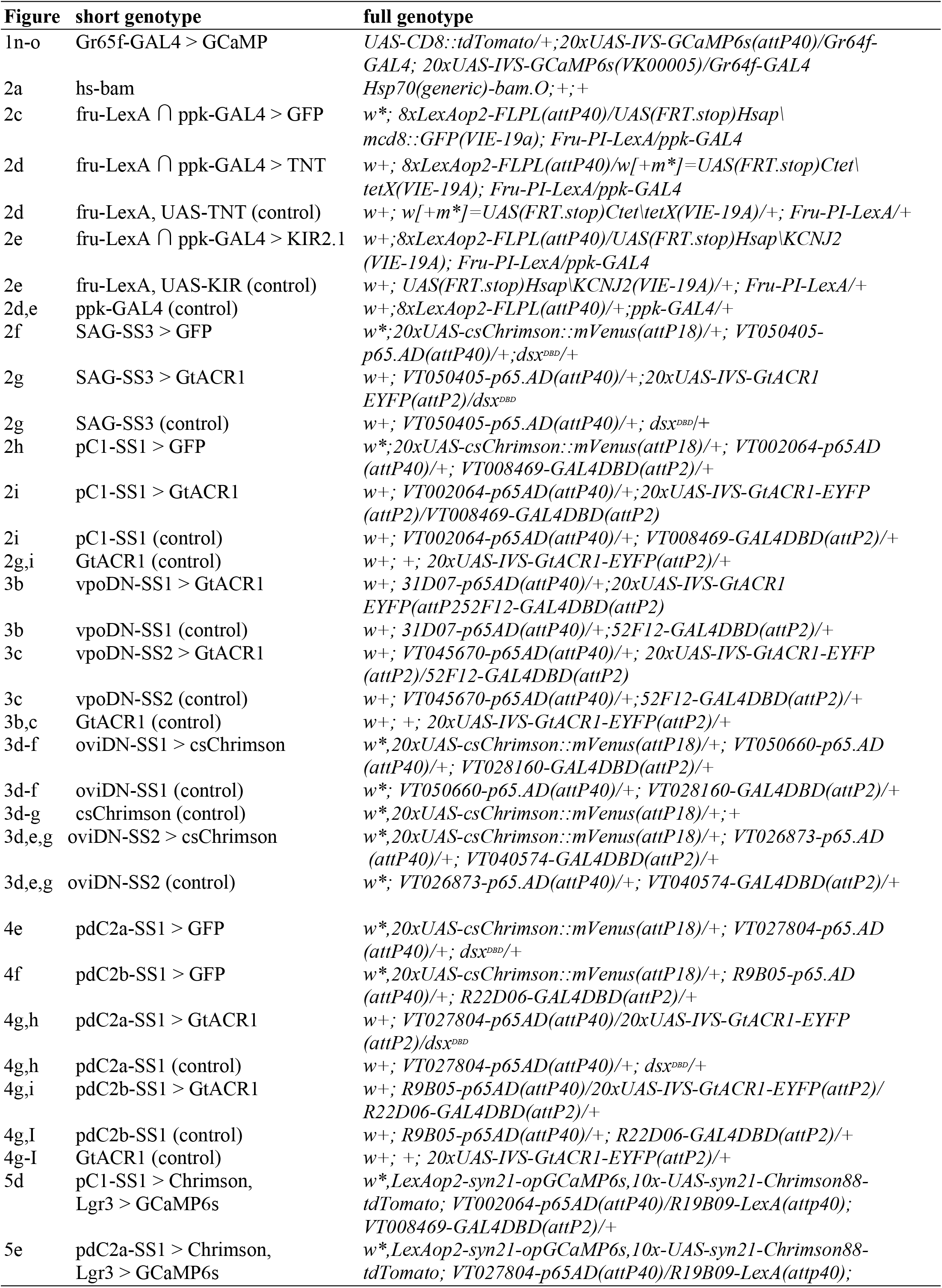
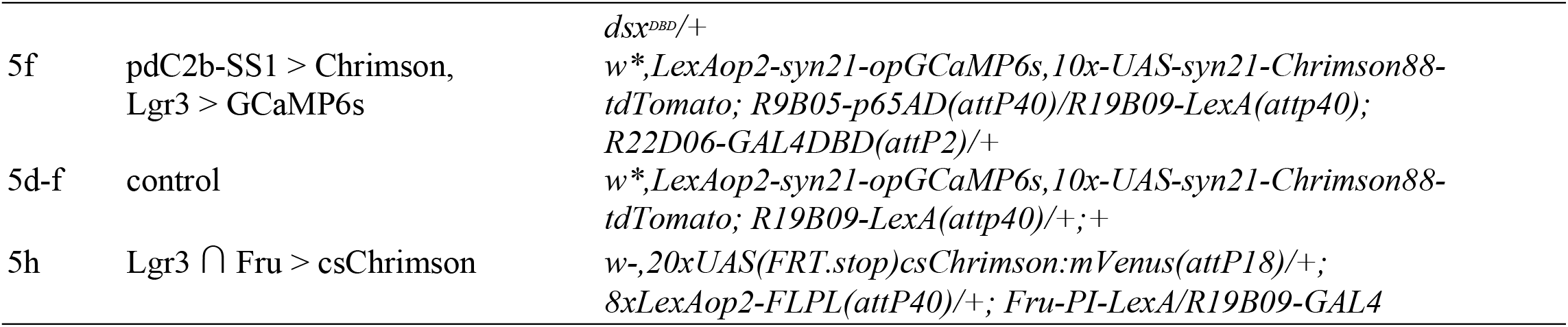
Fly genotypes in figures.

**Supplementary Table 2.**
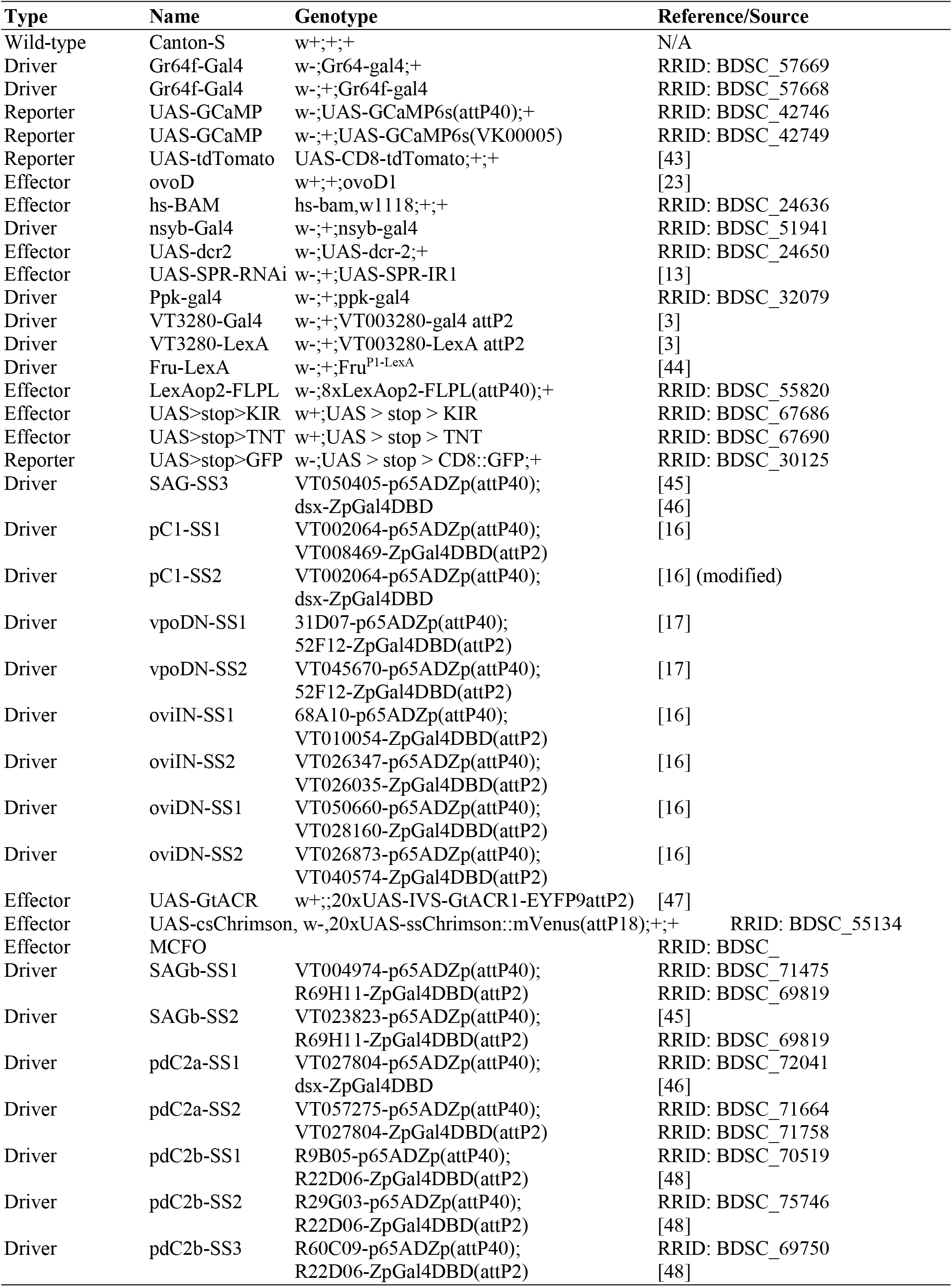
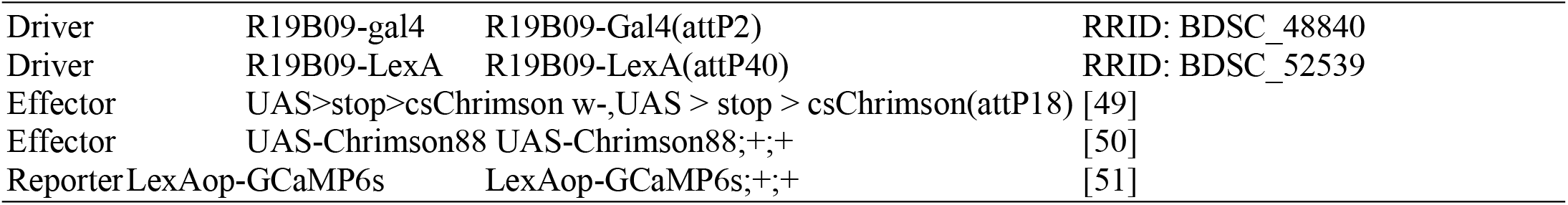
Fly Stocks.

## Acknowledgements

Members of the Scott lab provided contributions to experimental design, data analysis, and manuscript preparation. Salil Bidaye assisted with the analyses of locomotor data. This work was supported by NIH R01DC013280 (K.S.). Neuronal reconstruction for this project took place in a collaborative CATMAID environment in which 27 labs are participating to build connectomes for specific circuits. Therefore, work done for published (Wang et al. 2020) and ongoing projects has proved useful to us. Development and administration of the FAFB tracing environment and analysis tools were funded in part by National Institutes of Health BRAIN Initiative grant 1RF1MH120679-01 to Davi Bock and Greg Jefferis, with software development effort and administrative support provided by Tom Kazimiers (Kazmos GmbH) and Eric Perlman (Yikes LLC). We thank Peter Li for sharing his automatic segmentation (citation is https://doi.org/10.1101/605634). Tracing in Cambridge was supported by Wellcome Trust (203261/Z/16/Z) and ERC (649111) awards to G. Jefferis. We thank the Princeton FlyWire team and members of the Murthy and Seung labs for development and maintenance of FlyWire (supported by BRAIN initiative grant MH117815 to Murthy and Seung). We found the partial tracing of SAG and pC1 in CATMAID useful (Wang et al. 2020). In addition to our tracing efforts, we are appreciative of the tracing accomplished by other labs. For help with tracing of pC1 in FlyWire, we thank the Murthy and Seung labs (77% of pC1a, 98% of pC1b, 92% of pC1c) and the Jefferis lab (7% of pC1a, 8% of pC1c). For help with tracing pCd-2 cells we would like to thank the Dickson lab (CATMAID: 76% of pCd-2a, 44% of pCd-2b, 28% of pCd-2c), the Jefferis lab (CATMAID: 20% of pCd-2a, 6% of pCd-2b, 3% of pCd-2c; FlyWire: 32% of pCd-2a, 8% of pCd-2b, 13% of pCd-2c), the Murthy and Seung lab (FlyWire: 15% of pCd-2a, 11% of pCd-2b, 6% of pCd-2c), and Janelia tracers (FlyWire: 52% of pCd-2a, 14% of pCd-2b). We would like to thank the Jefferis labs (46%) and the Murthy and Seung labs (50%) for their contribution to tracing the t-shaped Lgr3 cells. The tracers who contributed include Nora Forknall, Lily Talley, Katie Stevens, Shanice Bailey, Tansy Yang, Istvan Taisz, Greg Jefferis, Lucas Encarnacion-Rivera, Austin T. Burke, Yijie Yin, James Hebditch, Kyle Patrick Willie, Joshua Bañez, Philipp Schlegel, Dharini Sapkal, Irene Salgarella, Dhwani Patel, Ben Silverman, Bhargavi Parmar, Chitra Nair, Doug Bland, Laia Serratosa, Marta Costa, Nash Hadjerol, Regine Salem, Rey Adrian Candilada, Shirleyjoy Serona, Zairene Lenizo, Zeba Vohra, and Zepeng Yao.

## Author contributions

M.L. and K.S. conceived and designed the study and wrote the manuscript. M.L. performed the experiments and analyzed the data with the exception of the functional imaging experiment, performed and analyzed by G.S.

## Corresponence and requests for materials

should be addressed to K.S. or M.L.

